# Detection of positive selection driving antimicrobial resistance in the core genome of *Staphylococcus epidermidis*

**DOI:** 10.1101/2024.09.30.615834

**Authors:** Callum O. Rimmer, Jonathan C. Thomas

## Abstract

*Staphylococcus epidermidis* is a commensal skin organism and leading cause of medical-device associated infections. Although previous research has investigated the phylogenetic diversity of the species, the level of positive selection on the core genome has yet to be explored. Here, we present the first core genome-wide screen of positive selection in the species. A curated dataset of 1003 whole-genome sequences (WGS) was created which represented the global diversity of the species, including all previously identified clades and genetic clusters (GCs). A 100-strain subset, which retained the diversity of the collection, was created by pruning the species-level tree with treemmer; core genes present in all genomes were extracted with Roary and used for positive selection analysis (n = 826). Site-level analysis was performed using PAML with omegaMap for confirmation. Selection along branches separating clades A and B were also investigated using PAML branch-site models and HyPhy. PAML site analysis revealed 17 genes under selection, including six hypothetical genes, most of which were linked to metabolism or transport. Several genes were associated with antimicrobial resistance, such as *ileS* which confers resistance to mupirocin. *cysG* and *sirC*, which catalyse the first two steps in the synthesis of siroheme, were also under selection. Two genes were found to be under selection at the branch-site level by both PAML and HyPhy, of which only one, *rhtC*, has been functionally characterised. Our analysis reveals the extent to which positive selection is operating on the core genome and identifies candidate genes which may have important roles in the fitness of the species.

## Introduction

*Staphylococcus epidermidis* is a Gram-positive opportunistic pathogen and member of the coagulase-negative group of staphylococci. An ubiquitous coloniser of the human skin, it is a frequent source of both contamination and infection, especially amongst hospital patients with indwelling medical devices such as catheters or central venous lines (1). The population structure of *S. epidermidis* has been extensively investigated (2–7), revealing a highly structured phylogeny composed of three major clades (genomic groups A-C) based on the core genome (8–10) while multilocus sequence types cluster into six different genetic clusters (GCs) (6,11). GCs one, five and six have been linked to pathogenic or generalist lifestyles and mostly correspond to genomic group A. STs belonging to GCs two and four (genomic group B) have been isolated from more niche environments, including sources such as rice grains and wild mice and often lack the genomic features linked to more pathogenic lifestyles. GC three (genomic group C) has been identified as a genetic sink with a large degree of admixture from the other clusters and appears highly recombinant (6,11). More recently, an alternative approach has been used whereby a score is attached to the collective sum of either accessory genes or SNPs in shared genes for a given strain, and has yielded improved accuracy for predicting isolation source (12). A recent pan-GWAS analysis supported previous findings that the two major clades of *S. epidermidis* are enriched for different genes, however it was also shown that these clades are associated with different body sites; isolates from clade B were enriched for feet sites whereas those from clade A showed no significant association with any particular location (13).

Positive selection is a process whereby mutations with a fitness advantage increase in frequency in the population, driven by selective pressures resulting in either directional, diversifying or balancing selection (14). Analysis of patterns of positive selection across the core genome may identify genes that play an important role in adapting to the environment. Since next-generation sequencing technologies have become more widely accessible, the number of publicly available genomes has grown exponentially. This has given more power to studies analysing selection as a larger sample of genomes, more representative of the overall population can be screened. When analysing selection most tools focus on the ratio of non-synonymous to synonymous mutations (ω) within genes, with the premise that a ω value of 1 represents neutral evolution, ω < 1 represents purifying selection and ω > 1 represents positive selection, particularly diversifying selection. While ω = 1 is regarded as the threshold for selection, it is rarely true for a gene to evolve strictly neutrally across the entire length of the gene, as evolution is typically constrained at functionally important sites within a gene. Hence the development of site models which allow the ω ratio to vary across a gene has been beneficial, as genes can now be flagged as under selection where previously the signal of selection would have been diluted since previous methods relied on an average ω value across the whole gene.

Identifying genes under selection using *in silico* approaches is well-established in multiple bacterial species (15–19), although in *S. epidermidis* while balancing selection has been investigated there has never been a genome-wide scan for positive selection (20). In this study we identify genes under positive selection across the core genome of a phylogenetically representative subset of the global *S. epidermidis* population. Specifically, we identify two sets of genes: 1) those where selection acted in any lineage throughout the species and 2) those that are under selection between the two major clades of *S. epidermidis* (genomics groups A and B).

## Methods

### Bacterial strains

We acquired 12 *S. epidermidis* strains from other laboratories whose STs belonged to the underrepresented GC2. Strains were cultured at 37°C overnight on trypticase-soy agar (Oxoid, UK; TSA). Single colonies were transferred to 10 mL trypticase-soy broth (Oxoid, UK; TSB) and incubated overnight at 37°C. Strains were stored at –80°C in a mixture of TSB and 15% glycerol (v/v). Genomic DNA was extracted with a Qiagen DNeasy kit according to manufacturer’s instructions (using 10 μL lysostaphin [1 mg/mL] for cell lysis).

### DNA sequencing and hybrid assembly

Illumina and Nanopore sequencing of *S. epidermidis* isolates was performed as described in Rimmer and Thomas, 2022 (21). All software was run according to default parameters unless otherwise noted. Fast5 sequencing reads were basecalled with the high accuracy model of Guppy v3.6.1 (Oxford Nanopore, UK). Sequence adapters were filtered using Porechop v0.2.4 (22) with middle and end thresholds of 85 and 95% respectively. Reads were filtered based on quality and length using Filtlong v0.2.1 (23). Canu v2.2 (24) was used to assemble overlapping reads into one contiguous sequence. The assembly was polished with four iterations of Racon v1.4.20 (25), followed by Medaka (-m r941_min_high_g360) v1.6 (26) and Nanopolish v0.13.2 (27). Trimmomatic v0.39 (28) was used to ensure all adapter sequences were removed from Illumina data. The output from Nanopolish (27) was error corrected with Illumina data using Racon and Pilon v1.24 (25,29). The assemblies were manually trimmed and re-orientated to *dnaA* using Circlator v1.5.5 (30). Assembly quality was determined using CheckM (31) and average nucleotide identity (ANI) was compared to the *S. epidermidis* type strain NCTC 11047^T^ using FastANI (32). All assembly metrics are available in supplementary table 1.

### Public whole-genome sequences

In October 2020 1,272 whole-genome sequences of *S. epidermidis* were downloaded from online repositories: 862 from NCBI (33), 72 from PubMLST (34), 97 from Dryad (35) and 241 from figshare (36). In May 2022 an additional 351 WGS were downloaded from NCBI (33) and 19 from PubMLST (34). This resulted in a collection of 1,642 WGSs before filtering. As described previously, ANI was calculated using FastANI with *S. epidermidis* type strain NCTC 11047^T^ as a reference. Assembly quality was determined by CheckM. Any genome with an ANI value of < 95% compared to the type strain was removed. Genomes with a size of less than 2.325 Mbp or greater than 2.9 Mbp, N50 less than 50 Kbp, more than 180 contigs, contained any ambiguous bases, contamination of more than 2.5% or completeness of less than 95% were removed. In total, 651 genomes were removed, leaving 991 WGS for analysis. The 12 hybrid assemblies sequenced in-house were added to the public dataset, leaving 1003 genomes to carry forward for analysis.

### Sequence typing and genetic cluster assignments

Sequence types (ST) were assigned using mlst (37) with the Thomas et al. *S. epidermidis* scheme (4). Novel loci were submitted to the PubMLST *S. epidermidis* database. Sequence types were assigned to GCs with BAPS v6 (38) using a codon linkage model. Allelic profiles and their alleles were downloaded for all 1,158 STs in PubMLST’s *S. epidermidis* database as of 24/06/2022. Allele sequences were trimmed and reverse complemented where necessary to maintain the +1 reading frame, before being concatenated into a single sequence. BAPS was run independently five times, with the maximum number of clusters set from 11 to 20. All ST and GC information is available in supplementary table 1.

### Determining the core genome and phylogenetic analysis

All 1003 genome assemblies were re-annotated with Prokka v1.14.6 (39) to ensure standardised gene annotations across the dataset. Roary v3.13 (40) was used to determine the core genome using gff files from Prokka. Flags ‘-e’ and ‘-n’ were used to create core gene alignments with MAFFT and flags ‘-cd 100’ and ‘-z’, defining core genes as those present in 100% of the dataset and keeping intermediate files, respectively. Roary identified 533 genes as core, however four were multi-copy genes and removed, leaving 529 core genes. The multiple sequence alignments (MSAs) for the 529 core genes were scanned for recombination using RDP5 v5.5 (41) using four recombination tests: RDP, GENECONV, Chimaera and MaxChi. The highest acceptable *P*-value was set to 0.05 and Bonferroni correction was applied. Genes where recombination events were detected by three or more tests were removed, leaving 467 recombination-free core gene MSAs for further analysis. All 467 MSAs were concatenated into a single sequence for all 1003 strains using FASconCAT v1.11 (42). A Maximum-likelihood tree was produced using RAxML-NG v1.1.0 (43) with tree model GTR+G+I and 100 bootstrap replicates and visualised with iTOL (44).

### Creating a reduced dataset for positive selection analysis

Positive selection analysis on gene alignments of 1003 strains was not computationally feasible; it was necessary to reduce the size of the dataset but still reflect the phylogenetic structure of the original tree. Treemmer v0.3 (45) was used to prune the 1003 strain phylogenetic tree down to 100 strains using ‘-X 100’ based on metadata containing the GC assignments, complete assemblies and strains sequenced in-house (− lm). The list of strains in the pruned tree was used as the reduced dataset. A new core genome was created with Roary as described above; both PRANK (46) and MAFFT were used for sequence alignments. In this reduced dataset, 1387 genes were identified as core. Core gene MSAs were checked for recombination with RDP5 as previously described; 381 genes were removed due to recombination, as high levels of recombination can result in a high rate of false positives (47). Alignments were scanned for gaps and stop codons with custom scripts; any genes with frameshift mutations in the MAFFT alignments were removed from the dataset, and terminal stop codons were removed using AliView (48). After curation, 826 genes were carried forward for analysis. Gene trees were generated using RAxML-NG with tree model GTR+G+I from the individual PRANK alignments.

### Positive selection analysis

For site-based analysis, the CODEML package within PAML v4.9 (49) was used to determine if any genes were under positive selection. Each gene was tested, using individual gene trees and the PRANK alignments, against two sets of site models: M1a vs M2a and M7 vs M8 (50,51). Each pair consists of a test model which allows selection and a null model where no selection is allowed. The log-likelihood scores (ℓ) for each were then compared using the likelihood-ratio test (LRT), calculated as 2Δℓ.

LRT values were converted to *P-*values using the chi2 program within PAML (df = 2) and the false discovery rate (52) was used for multiple correction testing. Three replicates were performed for each model, and genes were classed as under positive selection if the LRT was significant for each replicate. Positively selected sites were identified using the Bayes empirical Bayes test (53) within PAML. omegaMap was used as an independent confirmatory test for genes identified as under positive selection by the PAML site models (54). omegaMap is a phylogeny-free approach and only requires gene alignments. Variation in ω for each gene’s MSA was determined using the variable model, where each sequence was split into blocks of three codons; sites within each block share the same ω. Codon frequencies calculated by PAML were used for omegaMap. An inverse distribution of ω was used, with minimum and maximum values of 0.01 and 50, respectively. The Monte Carlo Markov chain (MCMC) was run for 250,000 iterations and 10 orderings.

The 100-strain subset core gene MSAs from PRANK were also concatenated into a single sequence and a species tree was generated with RAxML-NG with tree model GTR+G+I and 1000 bootstrap replicates, which was visualised with iTOL. To identify whether genes are under selection along the four branches between genomic groups A and B in the species tree (Figure 1), PAML branch-site models A and A_null_ were used (55,56). Five replicates were performed for each model using different initial values of ω (0.4 – 1.5). The highest log-likelihood score of the five replicates for each model was used to perform the LRT. LRT values were converted to *P-*values using the chi2 program from PAML (df = 1; FDR corrected). Given that omegaMap is a phylogeny-free approach, it could not be used as the independent test for selection on specific branches. Instead, MEME, BUSTED and aBSREL from the HyPhy package were used as alternative independent tests to identify genes under selection along the four branches (57–60). MEME was run with a *P*-value cut-off of 0.05 and 500 bootstraps. BUSTED and aBSREL were run with synonymous-rate variation (SRV). All three programs were run with ‘--kill-zero-lengths No’. FDR correction was not applied to the branch-site data due to lack of power when only testing four branches. As the computing time for HyPhy is considerably less than PAML, we also ran HyPhy on all core genes. PAML control files and the configuration files for omegaMap are available on GitHub (https://github.com/Callum-Rimmer/positive_selection). COG categories for all genes under selection were assigned using COGclassifier (61).

**Figure 1:**
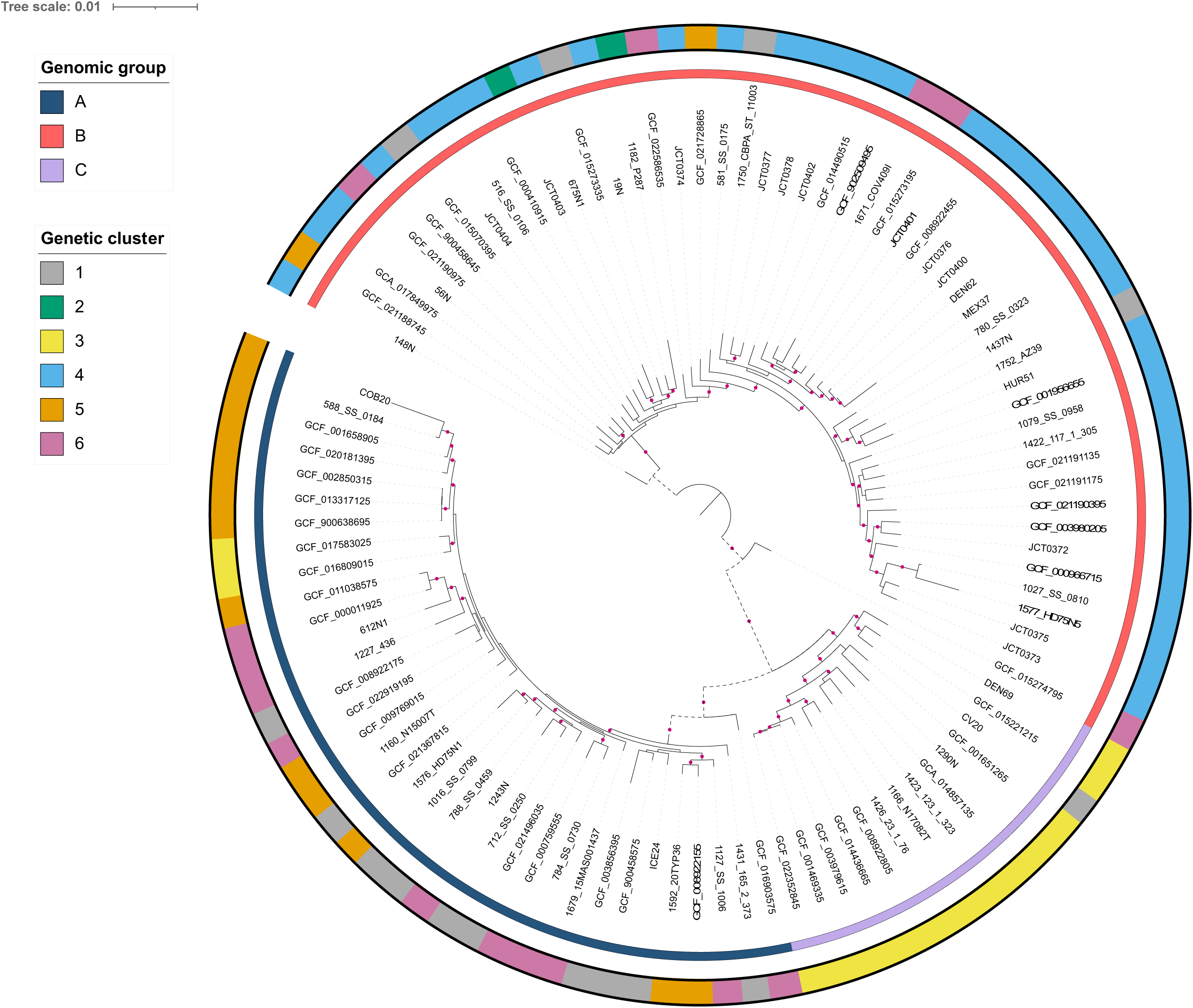
Maximum likelihood tree for the 100-strain subset, based on the concatenation of 826 core genes. Pink circles at nodes represent bootstrap support ≥ 75%. Inner and outer ring shows genomic group and genetic cluster, respectively. Branches with dashed lines were chosen for branch-site analysis with PAML and HyPhy.

## Results

### Curated database of WGSs

We created a final curated database of 1003 WGSs. The average genome size was 2.52 Mb, however there were significant differences between the genetic clusters (ANOVA P-value < 0.001). Genomes from GC5 were significantly larger compared to other clusters (on average 106 Kbp larger; Tukey HSD *P*-values < 0.001), except for GC2 which was not tested due to the small sample size. Genomes from GC4 were significantly larger compared to GC6 strains, with an average increase of 27 Kbp (Tukey HSD *P*-value = 0.006). Sixteen genomes were composed of a single contig. Excluding complete or nearly complete genomes (contigs < 10), the average N50 was 144 Kbp while the average number of contigs was 72. For the reduced dataset of 100 genomes, the average N50 was 427 Kbp while the average number of contigs was 64.

### MLST and genetic clusters

BAPS has previously been used to classify *S. epidermidis* sequence types into distinct genetic clusters. Since 2015 the *S. epidermidis* MLST database has expanded from 588 STs (11) to 1,158 STs, downloaded on 13/07/22. BAPS identified seven genetic clusters based on 1,158 ST profiles. Twenty-six STs that were previously assigned a GC by Tolo et al (11) were assigned a different GC with the new dataset although 24 of these involved GC3, a known genetic sink for the other clusters and subject to a large amount of admixture (11,62). Each of the five replicates produced identical results, however support for GC7 was limited with only seven STs. There was a diverse array of STs in our full curated dataset with 278 unique ST profiles. All genetic clusters except for GC7 were represented in the reduced dataset; GCs one, three, five and six each had 15 strains. GC2 had two strains and GC4 had 38 strains. This corresponded to 35 strains in genomic group A, 51 in group B and 14 in group C.

### Phylogenetic analysis

Supplemental figure 1 shows the *S. epidermidis* phylogeny using the curated dataset of 1003 WGSs from globally distributed strains. Our tree shows consistent clade structure compared to previous studies (8,10) with the majority of isolates clustering into genomic groups A and B. Genomic group A mostly comprised strains from GC1, GC5 and GC6; genomic group B mostly contained strains from GC4 and genomic group C predominantly corresponded to isolates from GC3. Strains belonging to GC2 also clustered with genomic group B as previously shown (8). No WGSs from GC seven were present in the dataset. While genomic group A was diverse with most strains belonging to three different GCs, there was clear separation of GC5 compared to GC1 and GC6. This is unsurprising considering strains from GC5 appear to be adapted to a hospital environment, while the majority of staphylococcal sequencing studies are focussed on clinical isolates (6,11). Figure 1 shows the phylogeny produced using the reduced dataset of 100 WGS; this tree shares the same clade structure as the complete 1003 strain dataset and shows the strains used for positive selection analysis are representative of the global population of *S. epidermidis*.

### Positive selection analysis

Nested site models from PAML identified 17 genes under positive selection (2.06% of core genes). All 17 were significant under the relaxed model set M7M8 while 10 were significant under the more stringent model set M1M2 (table 1). The Bayes empirical Bayes test embedded within PAML identified sites under selection for all genes except *serS*. Six hypothetical genes were under selection, five of which were significant with both model sets. omegaMap was also used to ensure that the list of genes under selection could be confirmed via an alternative independent test. Data from omegaMap agreed with 15 of the 17 genes flagged as under selection by PAML, with posterior probabilities of > 0.9 for each of the sites identified by the BEB test. omegaMap did not support selection for two of the genes, group_10327 and *serS* (no sites had a posterior probability of > 0.4). The BEB test from PAML did not identify any specific sites under selection from both M2 and M8 for *serS*, although it did for group_10327 (sites 335 and 341). While no COG categories were enriched for genes under selection, these genes belonged to a diverse set of classes, with 10 different COG categories across the 17 genes (nine COG categories for the 15 genes confirmed by omegaMap). Positions of genes under selection were determined in *S. epidermidis* strain RP62A and plotted to visualise. No putative hotspots for selection were identified.

**Table 1:** Genes under site-level selection based on PAML models M1M2 and M7M8. The Bayes empirical Bayes test from PAML model 8 was used to predict specific sites under selection. Hypothetical genes were denoted with the ‘group_’ prefix by Roary.

| Gene | M1M2 *p* value | Model 8 sites | M7M8 *p* value | RP62A locus tag | COG category | COG definition |
| --- | --- | --- | --- | --- | --- | --- |
| *csd* | <0.001 | 321 S** | <0.001 | SERP_RS02565 | E | Amino acid transport and metabolism |
| group_1172 | <0.001 | 66 G*; 160 R*; 202 S** | <0.001 | SERP_RS11475 | H | Coenzyme transport and metabolism |
| group_1349 | <0.001 | 183 Y*; 195 N**; 213 G**;<br>218 A*; 243 S** | <0.001 | SERP_RS04470 | V | Defense mechanisms |
| group_2443 | <0.001 | 151 H**; 156 G**; 214 P*;<br>552 L**; 584 V* | <0.001 | SERP_RS11155 | J | Translation, ribosomal structure and biogenesis |
| group_3177 | <0.001 | 463 S** | <0.001 | SERP_RS02575 | O | Posttranslational modification, protein turnover, chaperones |
| group_10276 | <0.001 | 107 K**; 167 D** | <0.001 | SERP_RS10320 | M | Cell wall/membrane/envelope biogenesis |
| *hisI* | <0.001 | 122 D** | <0.001 | SERP_RS11315 | E | Amino acid transport and metabolism |
| *ndhB* | <0.001 | 158 R*; 441 L** | <0.001 | SERP_RS00545 | C | Energy production and conversion |
| *sirC* | <0.001 | 54 A**; 89 G** | <0.001 | SERP_RS10840 | H | Coenzyme transport and metabolism |
| *yfcA* | <0.001 | 13 V **; 211 F** | <0.001 | SERP_RS06320 | P | Inorganic ion transport and metabolism |
| *cysG* | NS | 32 S**; 242 I**; 270 R** | <0.001 | SERP_RS09915 | H | Coenzyme transport and metabolism |
| group_10327<sup>α</sup> | NS | 335 H**; 341 Q* | <0.001 | SERP_RS12385 | T | Signal transduction mechanisms |
| *ileS* | NS | 588 V* | <0.001 | SERP_RS03840 | J | Translation, ribosomal structure and biogenesis |
| *nasD* | NS | 27 Q**; 757 V* | <0.001 | SERP_RS09925 | C | Energy production and conversion |
| *rluD_3* | NS | 77 V*; 174 A**; 179 L*; 201 K* | <0.001 | SERP_RS03005 | J | Translation, ribosomal structure and biogenesis |
| *serS*<sup>α</sup> | NS | NS | <0.001 | SERP_RS12455 | J | Translation, ribosomal structure and biogenesis |
| *yheS* | NS | 59 H*; 346 R*; 352 V* | <0.001 | SERP_RS08285 | R | General function prediction only |

\*Posterior probability > 95%, \*\*Posterior probability > 99%, NS Not significant
<sup>α</sup> Gene not supported by omegaMap

PAML branch-site models A/A_null_ identified 19 genes under selection (2.3% of core genes; table 2) along the branches between genomic groups A and B (highlighted as dashed branches in figure 1). The BEB test only identified sites under selection with a posterior probability of > 0.9 in four genes. Two genes were shared between the analyses of PAML branch-site models and HyPhy (both aBSREL and BUSTED): group_1158 and *rhtC*. For group_1158, PAML and MEME both reported codons 296 and 376 under selection (posterior probability > 0.91). MEME was able to link these two sites to foreground branches (one branch for site 296 and two branches for site 376) and reported a much higher LRT for site 376, similar to PAML. MEME also identified codons 68 and 196 as under episodic selection. group_1158 mapped to COG category U (intracellular trafficking, secretion and vesicular transport). For *rhtC*, both PAML and MEME predicted codon 94 under selection. MEME and aBSREL linked this site to one foreground branch. *rhtC* belongs to COG category E (amino acid transport and metabolism). While aBSREL and BUSTED did not identify selection in group_1315 and group_1893, MEME did support the sites highlighted by the BEB test. For HyPhy, aBSREL and BUSTED supported only two of the 19 genes identified by PAML: group_1158 and *rhtC*. MEME was able to identify sites under selection for 13 of the genes. After screening all core genes with HyPhy, signatures of selection were identified for 14 genes (one identified by only BUSTED; seven identified only by aBSREL, while six genes were significant with both). MEME was only able to identify specific sites under selection in 11 of the 14 genes; the three genes with no specific sites were not significant when testing with BUSTED (supplementary table two).

**Table 2:** Genes under branch-site level selection based on PAML models A / A_null_, with supporting data from HyPhy. aBSREL indicates how many of the four branches chosen to analyse are under selection, while BUSTED presents a single *P-*value for all branches. MEME presents a list of sites under selection similar to PAML.

| Gene | PAML<br>*p* value | PAML BEB sites | aBSREL:<br>branches under<br>selection | BUSTED<br>*p* value | MEME: sites under<br>selection | RP62A locus<br>tag | COG<br>category | COG definition |
| --- | --- | --- | --- | --- | --- | --- | --- | --- |
| *azoI* | 0.017 | - | NS | NS | NS | SERP_RS01225 | C | Energy production and conversion |
| *brnQ_2* | 0.016 | - | NS | NS | 40, 432, 433, 451 | SERP_RS11610 | E | Amino acid transport and metabolism |
| *femB* | 0.031 | - | NS | NS | 112, 147, 385 | SERP_RS04745 | M | Cell wall/membrane/envelope<br>biogenesis |
| *ftsZ* | 0.016 | - | NS | NS | 370 | SERP_RS03805 | D | Cell cycle control, cell division,<br>chromosome partitioning |
| group_1158 | 0.001 | 296, 376* | 1 | 0.004 | 68, 196, 296, 376 | SERP_RS06800 | U | Intracellular trafficking, secretion, and<br>vesicular transport |
| group_1315 | < 0.001 | 40**, 62*, 76** | NS | NS | 16, 40, 76 | SERP_RS00405 | - | - |
| group_1390 | 0.001 | - | NS | NS | 76, 89, 94 | SERP_RS10460 | - | - |
| group_1893 | 0.018 | 117* | NS | NS | 57, 117 | SERP_RS01795 | S | Function unknown |
| group_3008 | 0.003 | - | NS | NS | 69 | SERP_RS04320 | X | Mobilome: prophages, transposons |
| group_3726 | 0.041 | - | NS | NS | NS | SERP_RS00430 | - | - |
| group_9842 | 0.006 | - | NS | NS | NS | SERP_RS06520 | R | General function prediction only |
| group_10354 | 0.014 | - | NS | NS | NS | SERP_RS12430 | S | Function unknown |
| *gtfA_3* | < 0.001 | - | NS | NS | 2, 80, 167, 169, 258,<br>313, 315, 320, 323, 348 | SERP_RS01235 | M | Cell wall/membrane/envelope<br>biogenesis |
| *lysP_1* | < 0.001 | - | NS | NS | NS | SERP_RS04540 | E | Amino acid transport and metabolism |
| *purl* | 0.016 | - | NS | NS | 561, 615, 696, 722 | SERP_RS03330 | F | Nucleotide transport and metabolism |
| *recQ_2* | < 0.001 | - | NS | NS | 98, 163 | SERP_RS02045 | L | Replication, recombination and repair |
| *rhtC* | < 0.001 | 94 | 1 | 0.004 | 94 | SERP_RS00415 | E | Amino acid transport and metabolism |
| *yehR* | 0.026 | - | NS | NS | 11, 58, 66, 95 | SERP_RS10160 | S | Function unknown |
| *yjdJ* | 0.010 | - | NS | NS | NS | SERP_RS10350 | R | General function prediction only |

\*Posterior probability > 95%, \*\*Posterior probability > 99%, NS Not significant, BEB Bayes empirical Bayes test

### Distribution of mutations between genomic groups for sites under selection

After identifying specific sites under selection, we analysed the frequency of alleles at these sites in the original alignments from our 1003 strain dataset. For the 10 characterised genes under selection, most of the minor alleles (all alleles excluding the most frequently observed at a given site) in *ileS*, *rluD_3* and *yheS* could be attributed to genomic group A, at 97, 69 and 54% of all minor alleles, respectively (figure 2). While minor alleles were evenly distributed across genetic clusters for *rluD_3* and *yheS*, 90% of minor alleles at the selected site in *ileS* were from hospital-associated GC5 strains. Although only one site was under selection for *ileS*, for both *rluD_3* and *yheS* at least three sites showed evidence of selection. Despite most minor alleles belonging to genomic group A strains for these three genes, this was not consistent across all sites. For codon 179 in *rluD_3*, 75% of strains encoding a leucine were group B strains while at codon 346 in *yheS*, 97% encoding arginine were group B. For the remaining seven characterised genes, minor alleles predominantly belonged to genomic group B strains, ranging from 83-97% of all minor alleles. Only two selected sites had minor alleles that were primarily linked to strains belonging to Genomic group C; strains encoding valine at codon 242 in *cysG* and threonine at codon 13 in *yfcA*. As GCs one, five and six were well represented in our dataset as part of genomic group A, we wanted to investigate whether there was any differences in SNPs across these clusters. At codons 77 and 179 for *rluD_3*, over 80% of GC1 and GC6 strains encoded the major allele while roughly 78% of GC5 strains encoded the minor allele. This pattern was also seen at two codons for *yheS*.

**Figure 2:**
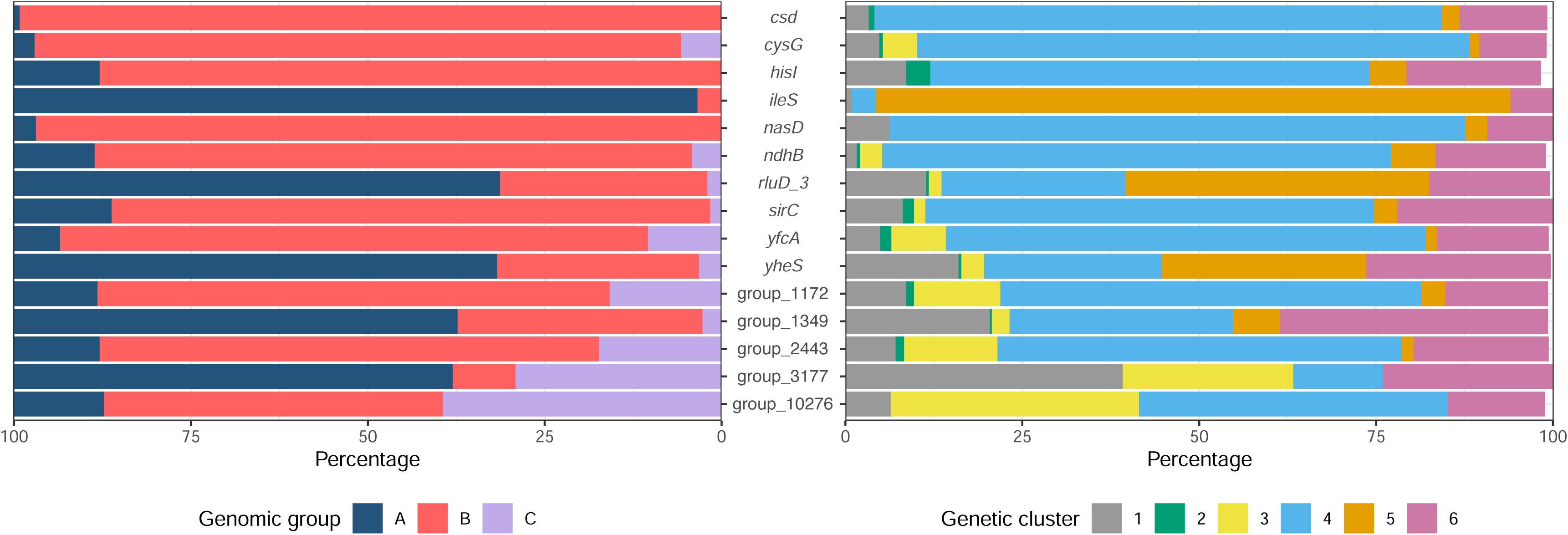
Distribution of minor alleles for each gene under site-level selection from PAML, split by genomic group (left) and genetic cluster (right). Analysis is based on the alignment data for all 1003 strains. Minor alleles are defined as all alleles excluding the most frequently observed at a given site.

For the five hypothetical genes under selection, minor alleles were largely observed in group A strains for group_1349 and group_3177, and group B strains for group_1172, group_2443 and group_10276. group_10276 also featured the most minor alleles attributed to genomic group C strains.

## Discussion

Identifying genes under positive selection is key to understanding how natural selection has shaped the evolution of an organism, both at the species level and within specific lineages of a species (63). Most studies that have performed genome-wide screens of positive selection typically use a small number of genomes, which makes it difficult to capture an accurate representation of the core genome of a species; this is largely due to either a lack of sequenced genomes and / or computational feasibility. Here, we have used 100 high-quality genomes that provide a balanced coverage of the three major clades and six genetic clusters of *S. epidermidis*.

Before the development of site models which allow for variation in ω across individual sites in a sequence, studies were mostly limited to gene-wide averages of the dN/dS ratio, meaning the strength of selection had to cover a substantial portion of the gene for it to be possible to sufficiently detect the signal to indicate a gene is under selection. While neutral theory plays a vital role in population genomics (64,65), multiple studies looking at signatures of selection within the core genome of bacterial species have observed extremely low average ω values: 0.16 in *Legionella pneumophila* (15), 0.05 in *Verminephrobacter* (16) and 0.064 in *Streptococcus dysgalactiae* (17). We observed an average ω value across all genes tested for selection of 0.12, indicating that most core genes within *S. epidermidis* are also under strong purifying selection; this includes genes which were themselves identified as under positive selection, for example group_1349 which had five sites under selection still had a gene-wide average ω value of 0.43. This highlights the importance of using site or branch-site models when analysing selection instead of gene-wide dN/dS values. There are exceptions to this, such as high sequence diversity, for example in studies screening for selection across multiple species (66–68).

Most genes linked to antimicrobial resistance or niche adaption in *S. epidermidis* were not investigated in this study as the majority of them are accessory genes. This pattern has been observed in multiple studies of other Gram-positive organisms: in *Staphylococcus aureus*, a genome-wide screen of 14 livestock-associated isolates found 60 genes under positive selection that were largely linked to metabolism (COG categories P, ion transport and metabolism; E, amino acid transport and metabolism; C, energy production and conversion) (69). In *Bacillus* species, studies have shown between 5 and 10% of the core genome is under positive selection, depending on the number of genomes screened; most genes under selection are linked to metabolism (18,70). Only 0.38% of core genes in *Pseudomonas aeruginosa* were under selection, and most genes were either proteases and hydrolases, transporters or associated with DNA stabilisation and replication (19).

## Selection at the site-level

We detected 17 genes under selection from PAML site models, approximately 2.1% of the core genome. While we could not perform COG enrichment analysis due to the diverse array of categories identified, most of the genes under selection are related to metabolism or are membrane associated. This is unsurprising given that membrane-associated proteins are the most likely to encounter selective pressures due to their contact with the environment (71).

Of the 17 genes PAML identified as under selection, two of them were not supported by omegaMap and a further five were hypothetical genes. The genes *cysG* and *sirC* catalyse the first two reactions in the three-step siroheme biosynthesis pathway (as a uroporphrinogen III methyltransferase [UroM] and precorrin-2 dehydrogenase [P2D], respectively) while *nasD* (*nirB*) encodes the large subunit of the nitrite reductase enzyme NirBD. The siroheme biosynthesis pathway was recently characterised in *S. aureus* and it was shown that the regulator of the *nir* cluster *nirR* is in fact a sirohydrochlorin ferrochelatase which catalyses the ferrochelation of sirohydrochlorin into siroheme (72). NirBD requires siroheme as a cofactor and converts nitrite (NO_3_^-^) into ammonia as a means of detoxifying the environment (73). It has been previously shown that high concentrations of nitrite derivatives can disrupt biofilm formation in *S. epidermidis* and mutagenesis studies of *sirC* showed manipulating a number of residues significantly increased the activity of the enzyme (73,74).

The genes *csd* and *yfcA* are both involved in the movement of sulphur between biomolecules (Csd is a cysteine desulferase while YfcA is a putative sulfur transporter belonging to the 4-Toluene Sulfonate Uptake Permease [TSUP] family). *csd* is involved in a number of different pathways, including the formation of iron-sulfur clusters and protection from oxidative stress and has recently been characterised in *S. aureus*; it is thought *csd*-like genes could be potential therapeutic targets (75,76). TSUP proteins are poorly characterised, however it was found TSUP homologues from *S. aureus* largely clustered as either Fe-S assembly proteins or transporters of sulfur-containing molecules (77). The fact that three genes directly linked to the siroheme biosynthesis pathway, an important cofactor for enzymes including NirBD, and a further two genes associated with the transport of sulphur and the formation of Fe-S clusters are under selection warrants further investigation.

*hisI* (*hisIE*) encodes a bifunctional enzyme which catalyses the second and third steps of the histidine biosynthesis pathway (78,79). Histidine biosynthesis plays an important role in bacterial metabolism however it has also been linked to pathogenesis; in *Acinetobacter baumannii* it was found extracellular histidine promoted pathogenesis while in *M. tuberculosis de novo* synthesis of histidine counteracted host upregulation of histidine catabolising enzymes (80,81).

Mutations in the *ileS* gene, which encodes an isoleucyl-tRNA synthetase (IleRS), have been extensively linked to mupirocin resistance in staphylococci (82–85). Mupirocin is a topical antibiotic typically used to decolonise patients of methicillin-resistant *S. aureus* (MRSA) by targeting bacterial IleRS (84). In *S. aureus*, it was found that a single residue change from valine to phenylalanine at codon 588 (which we have identified as under positive selection) resulted in low-level resistance to mupirocin and did not incur a large fitness cost (82). It is unsurprising widespread use of mupirocin has resulted in increased resistance in both *S. aureus* and *S. epidermidis* since they colonise similar body sites, however this highlights the need to look for alternative antimicrobial agents to decolonise MRSA or employ stricter antimicrobial stewardship. *rluD_3* appears to be a pseudouridine synthase responsible for converting uridine to pseudouridine in 23S rRNA; mutations in this gene have previously been associated with resistance to aminoglycosides (86–88). *ndhB* encodes one of the subunits of the type I NADH dehydrogenase, which forms part of the electron transport chain and is critical for bacterial metabolism (89,90). Mutations in NADH dehydrogenase genes have been linked to resistance against aminoglycosides due to the requirement of proton-motive force needed for uptake of the antibiotic (91). It has also been shown that mutations in *ndh* genes were linked to isoniazid resistance in *Mycobacterium tuberculosis*, where it is used as a treatment for both active and latent TB infections (92).

*yheS* encodes a putative ATP-binding cassette F (ABC-F) protein; these proteins are associated with antibiotic resistance, typically mediated through ribosomal protection where the ABC-F protein displaces the antibiotic at the ribosome binding site (93,94). ABC-F proteins are found in a wide array of bacterial species, with a number of them having been characterised in staphylococci (95–98).

There was a strong association between minor alleles and genomic group. For the three genes linked to AMR described above, most of these minor alleles were found in genomic group A. This is especially true for *ileS*, where the mutation was almost solely found in GC5 strains. This is unsurprising given previous work characterising strains from these genetic clusters (6,11), although this highlights the importance of monitoring antibiotic efficacy as we are observing strong evidence of a selection pressure amongst hospital-associated isolates. For the remaining genes, there was a clear association between the minor alleles and genomic group B / GC4. These genes are associated with key metabolic processes which are well understood, however it would appear that mutations in these genes are being driven by less evident selection pressures. Group B isolates includes both commensal strains and those from more unique environmental niches, however further study is needed to identify the role selection plays within this clade.

### Selection at the branch-site level

*rhtC* was the only characterised gene identified as under selection along the branches separating genomic groups A and B, from both the HyPhy and PAML branch-site models. Interestingly, while both the BEB test from PAML and MEME identified codon 94 under selection, the only observed change was a synonymous mutation of AGT to TCT (serine). This mutation was also almost exclusively observed in genomic group C strains, one group B strain also had this alternative codon but was the closest strain to group C on the tree. *rhtC* encodes a threonine efflux pump which has been shown to confer resistance to a variety of amino acid analogues (99,100). While the mutation in *rhtC* is synonymous, specific patterns of codon usage can impact both the speed and accuracy of translation and therefore affect fitness (101–103).

Using branch-site models to identify genes under selection along branches between genomic groups A and B was more challenging compared to the site models. With site models highlighting a few key sites under selection, repeating the same analysis using only four foreground branches posed a new challenge, since the proportion of sites under selection would be extremely low and so the signal present to be detected by PAML and HyPhy would also be much lower than during the site model tests. This was demonstrated by the BEB test in the PAML branch-site models; the BEB test was only able to highlight specific sites in four of the 19 genes under selection. Zhang et al. observed this issue in simulations of the PAML branch-site models; the power of the BEB test was low, and it frequently provided no sites under selection with a posterior probability of > 95% despite the branch-site model supporting selection on foreground branches (56). More recently Álvarez-Carretero et al. discussed how foreground branches in PAML branch-site models should be determined *a priori*, otherwise multiple correction testing is needed, while Bonferroni correction is too conservative for this type of analysis (63).

A similar issue is seen with branch-site models from HyPhy. Kosakovsky-Pond et al. described how the proportion of sites under selection required to detect episodic selection along foreground branches is very high, approximately 10-15% for a gene even with inflated ω values of at least four or five (104). However, when ω = 2, with the same proportion of sites under selection, the power of the branch-site model used dropped to 8%, which points to the fact that recognizing a weak selection signal is much more difficult, even with a high proportion of sites under selection (104). Therefore as the site model with PAML only identified a maximum of five sites under selection for any of the tested genes, given the low power of the branch-site model in the simulations described above, at the branch level it is difficult to detect a strong signal of selection. In addition, when testing datasets of viral pathogens using aBSREL, it was found that because of the large number of tests involved genes which showed significant support (uncorrected *P*-value < 0.05) for selection along foreground branches rarely survived multiple correction testing (59). This demonstrates that even with a large cohort of samples it is still challenging to reach the significance threshold with branch-level data, as shown with both our PAML and HyPhy results.

Unlike the PAML branch-site models, aBSREL and BUSTED do not explicitly test for selection at each site in a gene. aBSREL instead tests whether a proportion of sites for each of the test branches has evolved under positive selection while BUSTED tests for whether at least one site is under selection for at least one of the test branches. For the gene that is significant with BUSTED but not aBSREL, it is possible this is due to the strength of selection being too weak to flag as significant for a given branch with aBSREL, but across all four branches the collective signal is strong enough for BUSTED to detect selection. Conversely, seven genes were significant with aBSREL but not with BUSTED. It is likely the prior assumptions made by BUSTED, which fits a codon model with three rate classes did not fit as well compared to aBSREL, which infers the optimal number of rate classes for each branch, hence it was important to run both models.

## Limitations and conclusion

While isolates from North America and Europe are well represented in our dataset, there are few from Central and South America, Africa and Asia. As this work is based upon a ‘global’ phylogeny of *S. epidermidis*, important lineages could be missing, biasing the selection analysis towards western countries, despite featuring all of the major clades previously identified; in *E. coli,* lineages characterised as recombinant were later more accurately defined when using a larger, more diverse set of isolates (105). Insufficient isolates from GC2 were available to accurately capture information about the role selection has played in this cluster, although this is unsurprising given the rarity of isolates. Screening the entire 1003 isolate dataset for selection was not feasible due to the computing power required to run the analyses, particularly PAML. Even though this was mitigated by sub-setting the isolates based on the phylogeny and using the alignment data for the whole dataset when analysing allelic diversity, it is still possible that not all genes under positive selection were identified.

Understanding the level of positive selection across a core genome can reveal the selection pressures which have defined the evolution of a species. The candidate genes we have identified are typically related to core metabolic pathways or associated with antimicrobial resistance, which highlights that different selective pressures are driving natural selection in the three genomic groups within the population.

## Acknowledgements

We are grateful to Dr Ferran Navarro (Universitat Autònoma de Barcelona) for *S. epidermidis* strains belonging to genetic cluster 2.

## Author contributions

JCT contributed to conceptualization, methodology, project administration, resources, supervision and writing – reviewing and editing. COR contributed to data curation, formal analysis, investigation, methodology, validation, visualisation and writing – original draft.

## Data Availability

All data are available under BioProject accession number PRJNA1159912 and BioSample accession numbers SAMN43587831-SAMN43587842. Illumina raw read data and Nanopore base-called data were submitted to SRA and are available via accession numbers SRR30669856-SRR30669867 and SRR30669822-SRR30669833, respectively.

