## Supplementary material for "Detection of positive selection driving antimicrobial resistance in the core genome of *Staphylococcus epidermidis*": Supplemental legends.docx

Supplementary figure one: Difference in genome size between *S. epidermidis* genetic clusters.

Supplementary figure two: Maximum likelihood tree for the 1003 strain dataset, based on the concatenation of 467 core genes. Pink circles at nodes represent bootstrap support ≥ 75%. Inner and outer ring shows genomic group and genetic cluster, respectively. Branches with dashed lines were chosen for selection analysis.

Supplementary table one: Metadata for the genomes used in this study.

Supplementary table two: Genes under branch-site level selection based on screening all core genes from the 100 strain subset with HyPhy tools aBSREL, BUSTED and MEME.
