## Supplementary material for "Detection of positive selection driving antimicrobial resistance in the core genome of *Staphylococcus epidermidis*": Supplementary Table 2 - HyPhy only genes.docx

| Gene | aBSREL: branches under selection | BUSTED  *p* value | MEME: sites under selection | RP62A locus tag | COG category | COG definition |
| --- | --- | --- | --- | --- | --- | --- |
| *atpF* | 1 | NS | NS | SERP_RS08575 | C | Energy production and conversion |
| group_1158*^α^* | 1 | 0.001 | 68, 196, 296, 376 | SERP_RS06800 | U | Intracellular trafficking, secretion, and vesicular transport |
| group_1429 | 1 | NS | NS | SERP_RS08475 | - | - |
| group_3287 | 1 | NS | 22, 31, 63, 78 | SERP_RS08590 | - | - |
| *metK* | 1 | 0.009 | 162, 391 | SERP_RS06670 | H | Coenzyme transport and metabolism |
| *plsC* | 1 | NS | 68, 203 | SERP_RS06380 | I | Lipid transport and metabolism |
| *rhtC^α^* | 1 | 0.003 | 94 | SERP_RS00415 | E | Amino acid transport and metabolism |
| *ribE* | 1 | 0.022 | 155, 202 | SERP_RS06555 | H | Coenzyme transport and metabolism |
| *rpsQ* | 1 | NS | 23 | SERP_RS09145 | J | Translation, ribosomal structure and biogenesis |
| *sigB* | 1 | 0.026 | 30 | SERP_RS08395 | K | Transcription |
| *tal* | 1 | 0.007 | 50, 52, 162 | SERP_RS06595 | G | Carbohydrate transport and metabolism |
| *yedJ* | 1 | NS | 43, 157 | SERP_RS08490 | J | Translation, ribosomal structure and biogenesis |
| *yghA* | 1 | NS | NS | SERP_RS09595 | I | Lipid transport and metabolism |
| *yojF* | NS | 0.005 | 92 | SERP_RS01265 | - | - |

NS Not significant

^α^Supported by PAML
