## Supplementary figures and images for "Detection of positive selection driving antimicrobial resistance in the core genome of *Staphylococcus epidermidis*"

### Supplementary Figure 1 - Genome size plot.pdf

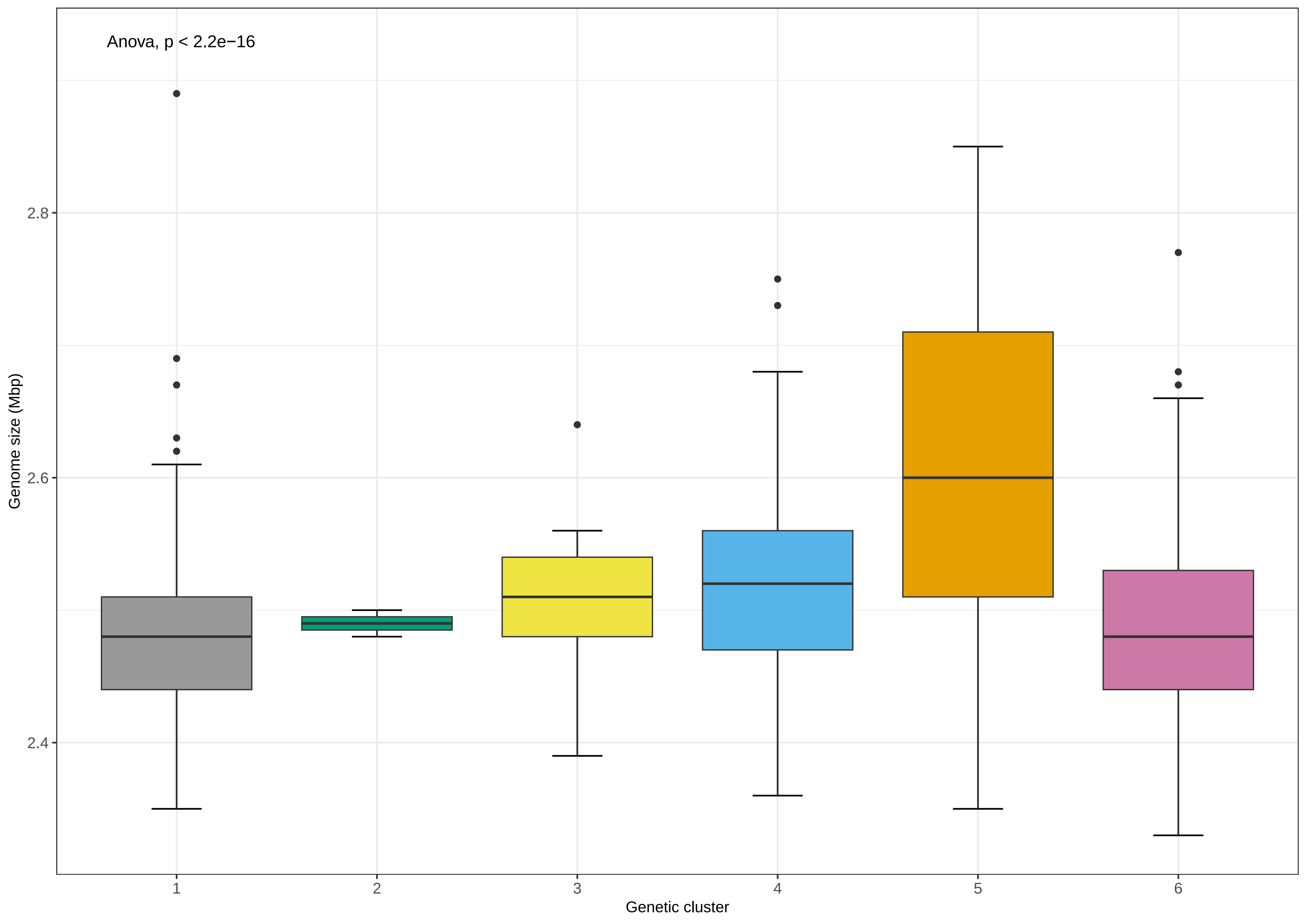

### Supplementary Figure 2 - S. epidermidis phylogeny of 1003 strains.pdf

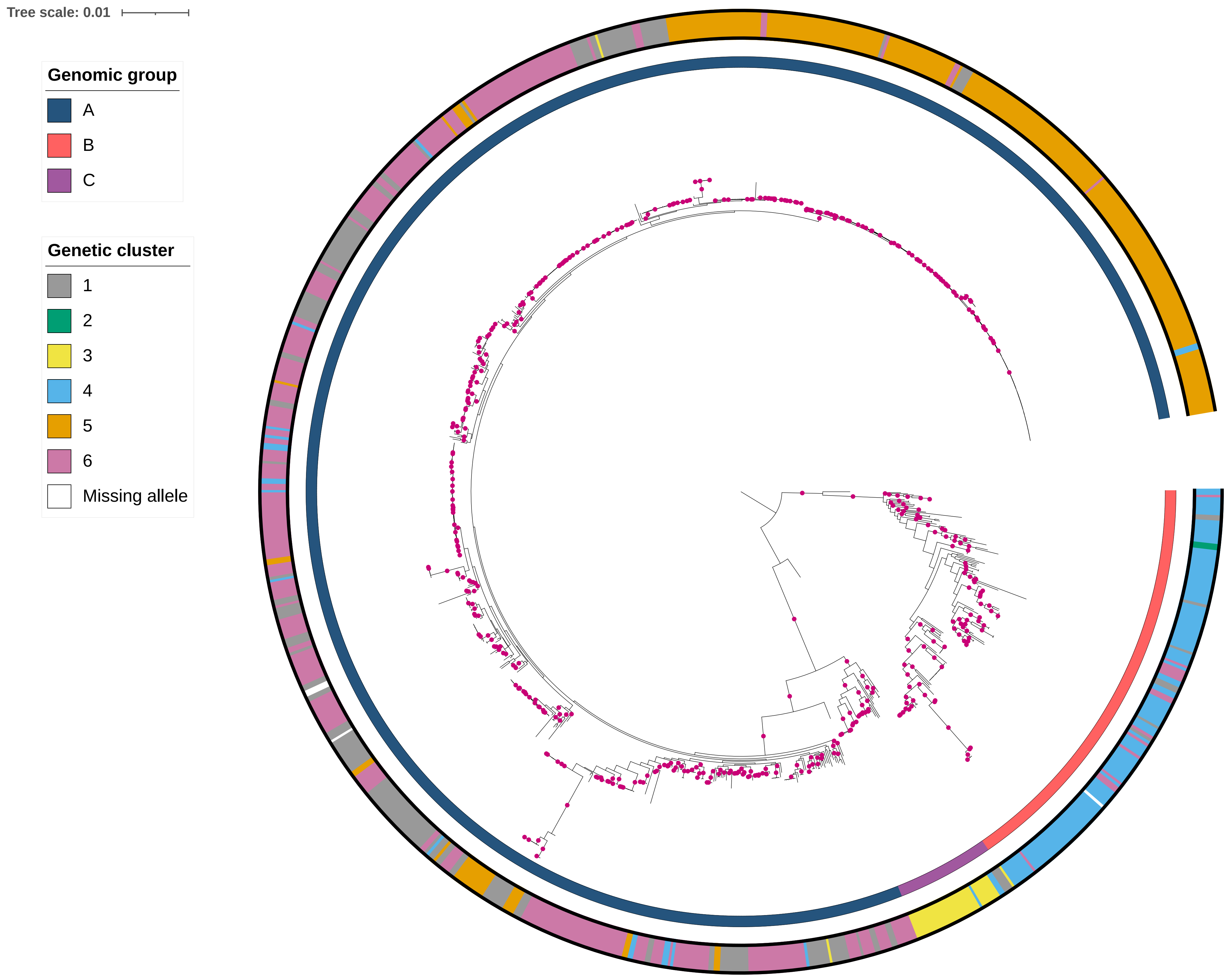
